# Molecular genomic studies of the obesogenic effects of tributyltin during adipogenic differentiation implicate a primary role for cytoskeletal damage

**DOI:** 10.1101/2022.06.28.497908

**Authors:** Taylor V. Thompson, John M. Greally

**Affiliations:** Center for Epigenomics and Department of Genetics, Albert Einstein College of Medicine, Bronx, NY, 10461, USA

## Abstract

Environmental obesogens are being studied for their potential role in the increasing prevalence of obesity globally. A major focus in this field of research has been on the mechanism by which these agents act. In this study we focused on the obesogenic organotin tributyltin (TBT), which is believed to act by binding to the PPARγ nuclear receptor in a heterodimer with RXR to alter gene regulation. To test whether this was the dominant mechanism for TBT activity, we performed time-course studies of transcription and chromatin accessibility in mesenchymal stem cells differentiating to adipocytes. We found limited evidence for PPARγ effects by TBT, but a strong response by Ras-related GTPases and evidence for the loss of TEAD transcription factor activity during differentiation. These observations combine to implicate a known property of organotins, to cause cytoskeletal cytoskeletal damage as the primary event in an updated model for TBT effects, leading to the loss of YAP co-regulator activity and the consequent failure of TEAD repression of adipogenesis.

## INTRODUCTION

While excess weight is typically believed to involve a caloric intake that is not matched by energy expenditure through exercise, the role of environmental obesogens is coming under increased scrutiny. The interest in this subclass of endocrine-disrupting chemicals is prompted in part by intriguing observations such as the increase over the last several decades of the average mid-life body weights in primates and rodents in research colonies, domestic dogs and cats, and feral rodents^1^. This observation occurs in parallel to the recognition that man-made chemicals that have ended up polluting the environment include biologically-active agents described as endocrine-disrupting chemicals^2^. Within this group exist chemicals whose effects involve promoting obesity, referred to as environmental obesogens^3, 4^. The paradigmatic environmental obesogen is the organotin tributyltin^5^, used in the production of plastics and anti-fouling paint on marine vessels, leading to the contamination of water systems and its exposure to humans through diet^6^.

Tributyltin (TBT) has been the focus of most experimental studies of environmental obesogens. A breakthrough set of observations in 2004-2006 was that TBT appeared to exert its effects through two transcription factors, the peroxisome proliferator-activated receptor gamma (PPARγ) and retinoic X receptor (RXR)^7–9^. This is an appealing model, as PPARγ is the master regulator of adipogenesis^10^, RXR is another nuclear receptor with which PPARγ forms a complex^11^, and inhibitors of both of these transcription factors (TFs) could reverse diet-induced obesity in human research participants^12^. The modelling of a crystal structure of the PPARγ ligand-binding domain with TBT^13^ added further weight to the evidence supporting TBT acting as a ligand of PPARγ to activate this nuclear receptor and promote adipogenesis. The possibility that exposure of TBT leads to inheritance of an obesity phenotype in subsequent non-exposed generations has also been a recent area of research activity^14, 15^.

We were interested in the potential for genome-wide studies of gene expression and its regulatory processes to understand how TBT exerts its adipogenic effects. With a focus on what we have described as cellular epigenetic models^16^, we wanted to explore how TBT exposure during early cell fate decisions could lead to cellular reprogramming, the alteration of characteristics of a canonical cell type. Molecular genomic assays that survey gene expression and chromatin organization would be expected to reveal PPARγ:RXR effects on the genome mediating their adipogenic effects at the cellular level, but there has been a surprising lack of such studies to date. The potential for these studies is to reveal the mediators at the genomic level of TBT effects, and to infer the upstream cell signaling driving these genomic regulatory changes. By testing human mesenchymal stem cells (MSCs), the effects at the earliest stages of adipogenic lineage commitment could be identified.

We chose a system of human MSC *in vitro* differentiation to adipocytes, exposing the cells throughout their differentiation to TBT and separately to the PPARγ agonist Rosiglitazone. By performing transcriptional and chromatin accessibility studies at four timepoints, we gained insights into the molecular genomic properties of cells undergoing normal adipogenic differentiation, and how the patterns deviate with the chemical exposures. The results indicate that TBT’s effects may be at least in part mediated by a PPARγ:RXR-independent effect on the cell, a novel insight into how some environmental obesogens may exert their damaging effects.

## RESULTS

### Characterization of the mesenchymal stem cell line

For our experiments we used the ASC52telo mesenchymal stem cell (MSC) line (ATCC SCRC-4000). This hTERT-transformed, adipose-derived cell line was first described in 2009^17^. The authors showed that these cells were able to differentiate to adipogenic and osteogenic lineages, and to have a distinctive pattern of surface antigen expression with flow cytometry. To test whether they met the consensus criteria to allow them to be described as MSCs^18^, we confirmed that the cells can be grown on a plastic surface, they express CD73, CD90 and CD105, but not CD45 (**Supplementary Fig. 1a**), and they can be differentiated *in vitro* to osteoblasts, adipocytes and chondroblasts (**Supplementary Fig. 1b**). The ASC52telo line therefore has the properties required to allow these cells to be described as MSCs.

### Cellular phenotyping following chemical exposures

We show an overview of the cellular phenotyping studies in **Fig. 1a**. We exposed the cells during *in vitro* differentiation to adipocytes to two chemicals. Our primary interest was in the effects of exposure to tributyltin (TBT), an environmental obesogen that acts as an endocrine-disrupting chemical^5^. Because of the prior studies implicating the mediation of TBT effects by PPARγ and RXR signalling^7–9^, we added the PPARγ agonist Rosiglitazone as a control exposure to represent the specific effects mediated by this nuclear receptor. As both agents were dissolved in dimethyl sulphoxide (DMSO), the equivalent amount was used as a vehicle control in the otherwise unexposed differentiating cells. The concentrations of TBT and Rosiglitazone used were prompted by those used in prior publications testing adipogenesis in comparable cell types^19, 20^.

**Figure 1:**
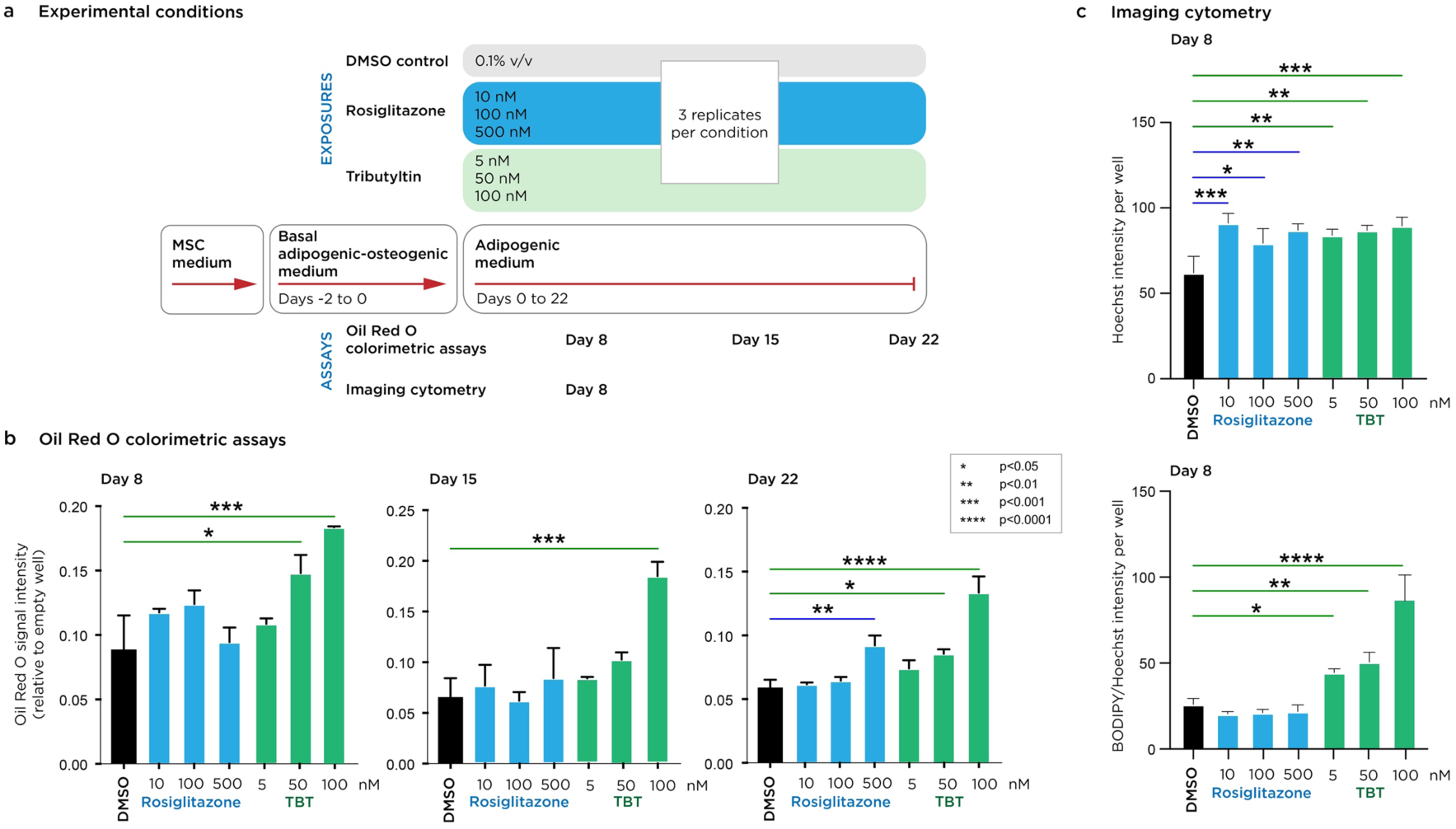
Cellular effects of Tributyltin (TBT) and Rosiglitazone. We show our experimental design in (a), sampling cells exposed to different concentrations of TBT and Rosiglitazone at multiple time points during adipogenesis. In (b) we show that TBT (green) leads to increased amounts of neutral lipid from as early as day 8, while Rosiglitazone (blue) begins to show increases by day 22. In (c) we show the results of studies that simultaneously quantify the amount of neutral lipid and cell numbers, showing that both TBT and Rosiglitazone cause cellular hyperplasia at all time points and doses, but that the neutral lipid content per cell is increased only for TBT.

We show in **Fig. 1b** how both TBT and Rosiglitazone increase the neutral lipid production by the cultured cells. An Oil Red O colorimetric assay revealed increased neutral lipid accumulation at the higher TBT doses by day 8 of differentiation, sustained at the highest (100 nM) dose through day 22, at which time the highest Rosiglitazone dose (500 nM) began to show significantly increased neutral lipid production (**Fig. 1b**). As the colorimetric assay does not distinguish between increased numbers of cells with no change in lipid content and increased amount of lipid per cell in the same number of cells, we used the Celigo system (Nexcelom) to image each cell using Bodipy to quantify neutral lipid content, and the Hoechst nuclear stain to count the number of cells. We focused on day 8 of differentiation when the earliest TBT effects were apparent. By this stage of differentiation both TBT and Rosiglitazone were causing cellular hyperplasia at all doses, as determined by Hoechst nuclear staining, but the dose-dependent accumulation of neutral lipid was limited to the TBT exposure (**Fig. 1c**). We conclude that both TBT and Rosiglitazone induce cellular hyperplasia, but that TBT induces neutral lipid accumulation at early stages of differentiation, with Rosiglitazone’s effects more limited and later in differentiation. The primary data from the imaging cytometry studies are provided in **Supplementary Table 1**.

### Transcriptional and chromatin studies of *in vitro* adipogenesis

To create a baseline against which to compare the effects of Rosiglitazone and TBT exposures, we repeated the adipogenic *in vitro* differentiation and sampled five timepoints, starting with cells in the basal adipogenic-osteogenic medium (Day 0) and then days 3 through 14 in adipogenic medium (**Fig. 2a**). RNA-seq and ATAC-seq^21^ were performed in triplicate at each of the 5 time points. The most substantial change in gene expression occurred between days 0 to day 3, following the transition to the adipogenic growth conditions (**Fig. 2b**), with a progression of more subtle transcriptional changes for the remainder of the differentiation. We studied panels of genes known to characterize white, beige and brown adipose tissue^22–24^, comparing their expression levels at the most differentiated stage (day 14) with the day 0 undifferentiated cells. In **Fig. 2c** we show how the genes characterizing white adipose tissue are the most consistently upregulated, indicating that the *in vitro* differentiation conditions of this MSC line generates cells with the properties of white adipose tissue.

**Figure 2:**
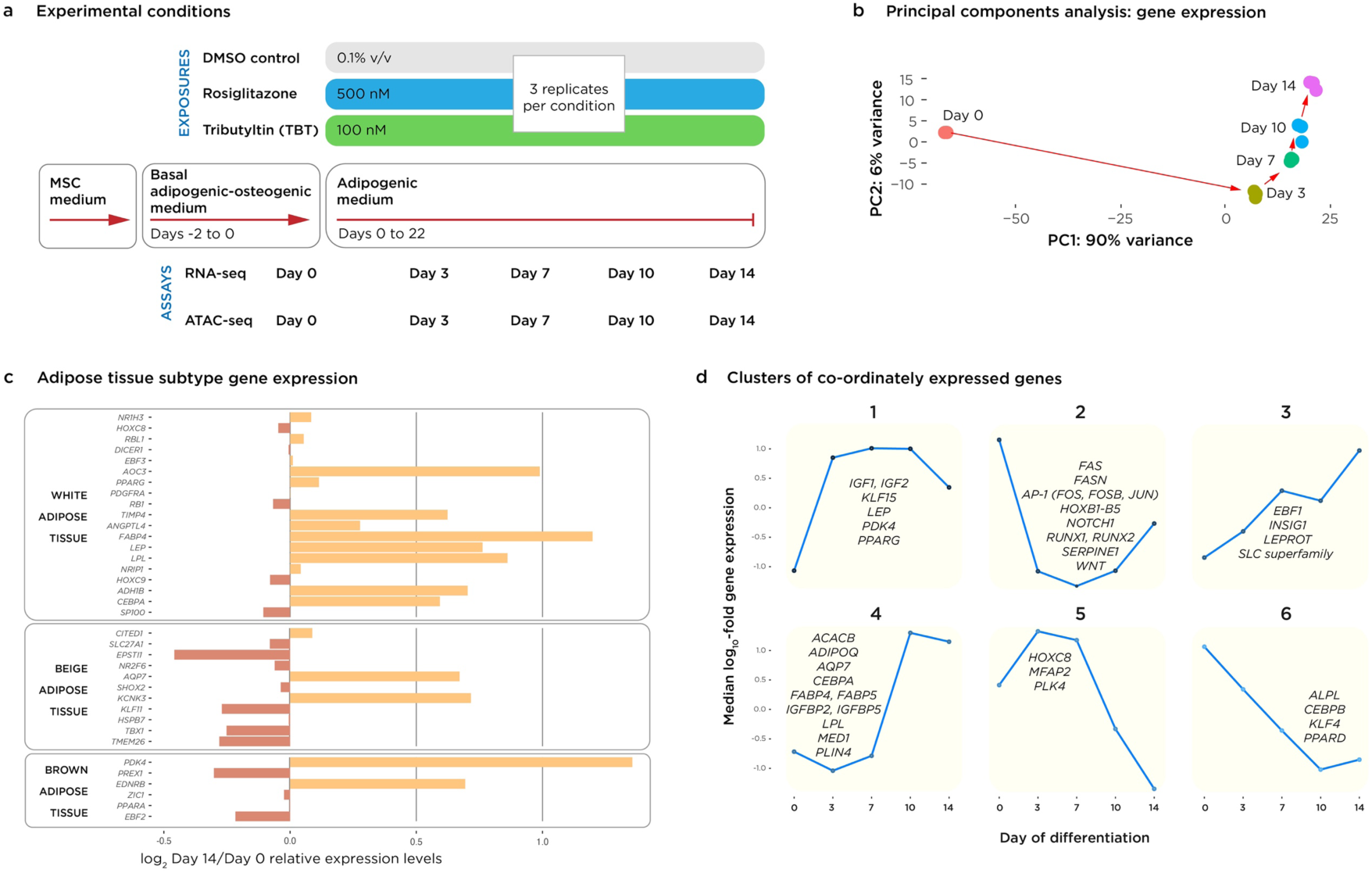
Molecular genomic studies of adipogenic differentiation. The choices of conditions for these experiments are shown in (a). Here we focus on the unexposed cells, using the same amount of the DMSO vehicle as was used to dissolve the TBT and Rosiglitazone. The most substantial change in gene expression occurs with the transfer to adipogenic medium (b), with continued, smaller degree changes thereafter. Using curated gene lists we show the relative expression of genes at day 14 compared to day 0 in (c), demonstrating the genes typical of white adipose tissue to be the most uniformly upregulated. In panel (d) we show examples of genes associated with each of 6 k-means clusters of distinctive patterns of expression over the time-course.

We used k-means clustering to study the temporal progression of expression of different groups of genes during adipogenesis. We illustrate 6 clusters in **Fig. 2d**. With the insight from **Fig. 2b** that the days 0 to 3 changes in gene expression are the greatest, we describe this as the ‘commitment’ stage, with days 3-7 the ‘consolidation’ stage and days 10-14 the ‘maturation’ phase. Clusters 1-2 reflect changes at the commitment stage, involving upregulation of genes encoding transcription factors (TFs) like *KLF15* and *PPARG* (cluster 1) and downregulation of other transcriptional regulatory genes, including multiple AP-1 and HOXB gene family members, *RUNX1* and *RUNX2* (cluster 2), and other transcriptional changes reflective of the acquisition of new metabolic properties by these cells. The consolidation stage is best represented by clusters 3 and 6, involving the induction of the *EBF1* (cluster 3) and down-regulation of the *CEBPB*, *KLF4* and *PPARD* (cluster 6) transcription factor genes. The maturation-associated clusters are 4-5, involving a further wave of TF expression changes, with upregulation of *CEBPA* and downregulation of *HOXC8* and *MFAP2*. These findings are consistent with and extend prior insights into the TF regulatory cascade during adipogenesis^25^. A list of all of the genes associated with each cluster is provided in **Supplementary Table 2**.

To understand how changes in expression of TF-encoding genes was reflected by their binding to regulatory loci in the genome, we performed ATAC-seq of the cells at the same time points. The loci that gained or lost transposase accessibility at each time point were identified and explored for TF-binding motifs. In **Fig. 3** we show representative results from this analysis (with complete results available in **Supplementary Table 3**). The AP-1 family of TFs is prominently represented at loci opening chromatin, reflecting its ability to recruit chromatin remodeling enzymes to specific genomic loci^26^. A number of the binding site motifs correspond to TFs known to be involved in adipogenesis, including the CEBP^27^, RUNX^28^, TEAD^29^, GATA^30^, EBF^31, 32^, NF1^33^ families of TFs, the ATF4^34^ TF and the CTCF^35, 36^ DNA-binding protein. These patterns of over-representation of TF binding sites provide us with a reference in normal differentiation against which to compare chemical-induced changes. We note that altered levels of expression of genes encoding TFs are not necessarily reflected by their regulatory activities, with the downregulation of AP-1 family genes following high expression levels at Day 0 in contrast to their ongoing inferred role in dynamic chromatin changes, illustrating the value of chromatin studies in addition to gene expression studies alone in understanding transcriptional regulatory processes.

**Figure 3:**
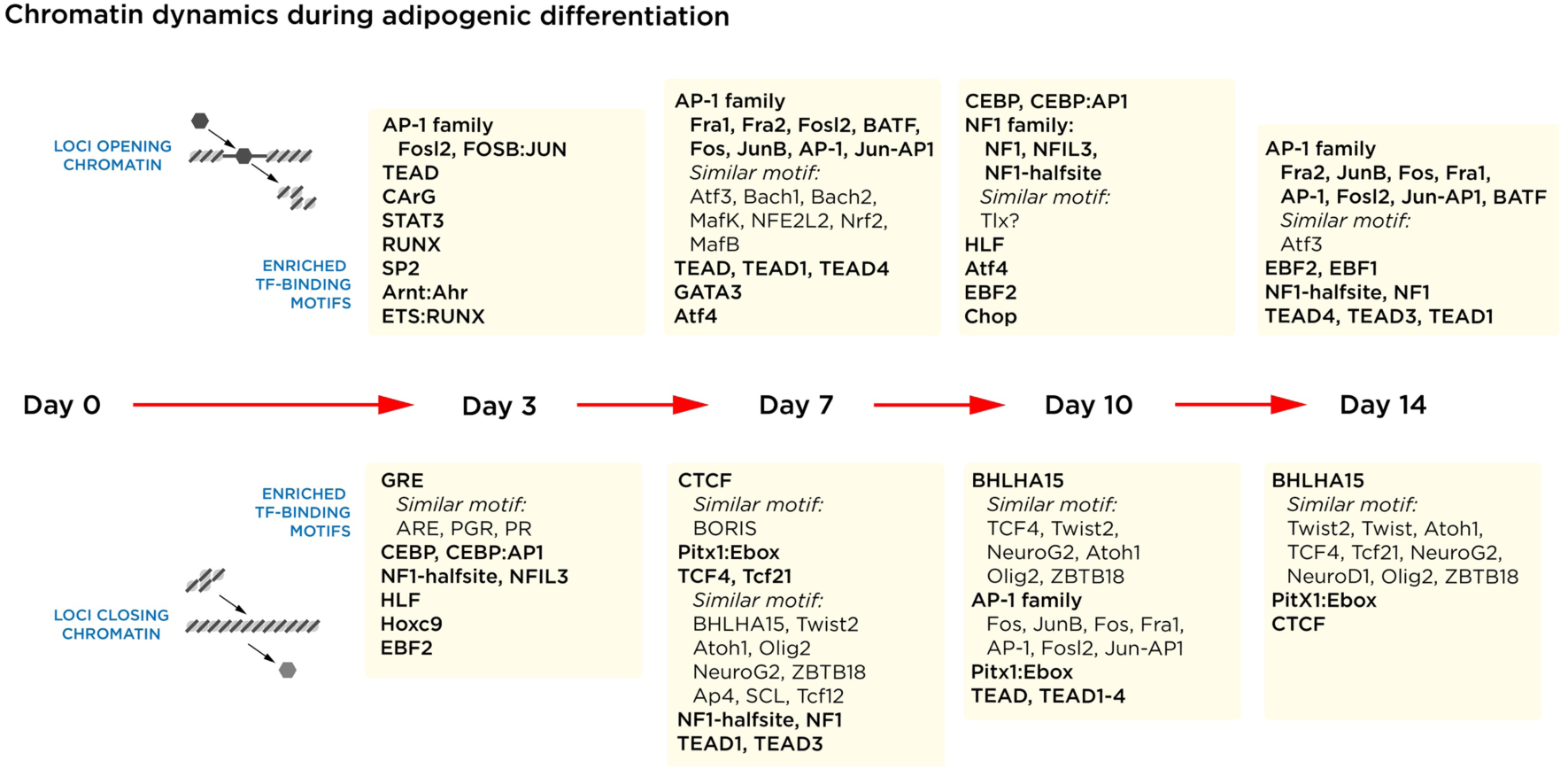
Inferred TF activities influencing chromatin organization during adipogenesis. By studying the DNA sequence motifs at the loci of accessible chromatin distinguishing each timepoint, we could infer the identity of the TFs at these loci. The results for each day represent the comparison with the previous timepoint, so those at day 3 represent loci gaining (upper) or losing (lower) accessibility compared with day 0. The AP-1 TF family has the capacity to recruit chromatin remodeling complexes to displace histones, making its presence in the loci gaining accessibility unsurprising. Many of these inferred TFs have known roles in adipogenesis, increasing our confidence in the validity of these results.

### Limited effects from exposure of adipogenic cells to the PPARγ agonist Rosiglitazone

Our cellular studies in **Fig. 1** showed that the PPARγ agonist Rosiglitazone induced cell proliferation by day 8 of *in vitro* adipogenesis, and increased neutral lipid accumulation by day 22. We used the experimental design of **Fig. 2a** and performed RNA-seq and ATAC-seq to characterize the molecular genomic effects of Rosiglitazone exposure early in adipogenic differentiation. We show the results in **Fig. 4**. The effects on gene expression were minimal, with few changes at any stage of the time course (**Fig. 4a**). The complete list of differentially expressed genes associated with Rosiglitazone exposure is provided in **Supplementary Table 4**. The patterns of gene expression for white, beige and brown adipose tissue were indistinguishable from the DMSO control; the cells were still producing white adipose tissue (**Fig. 4b**). When gene expression was compared for each of the days of differentiation, no significant differences were found for days 3 or 7, and limited changes at days 10 and 14. When the genes consistently increased in expression for those two time points were analyzed, a ChEA analysis^37^ performed using Enrichr^38^ showed evidence for PPARγ binding in proximity to the upregulated genes (**Fig. 4c**).

**Figure 4:**
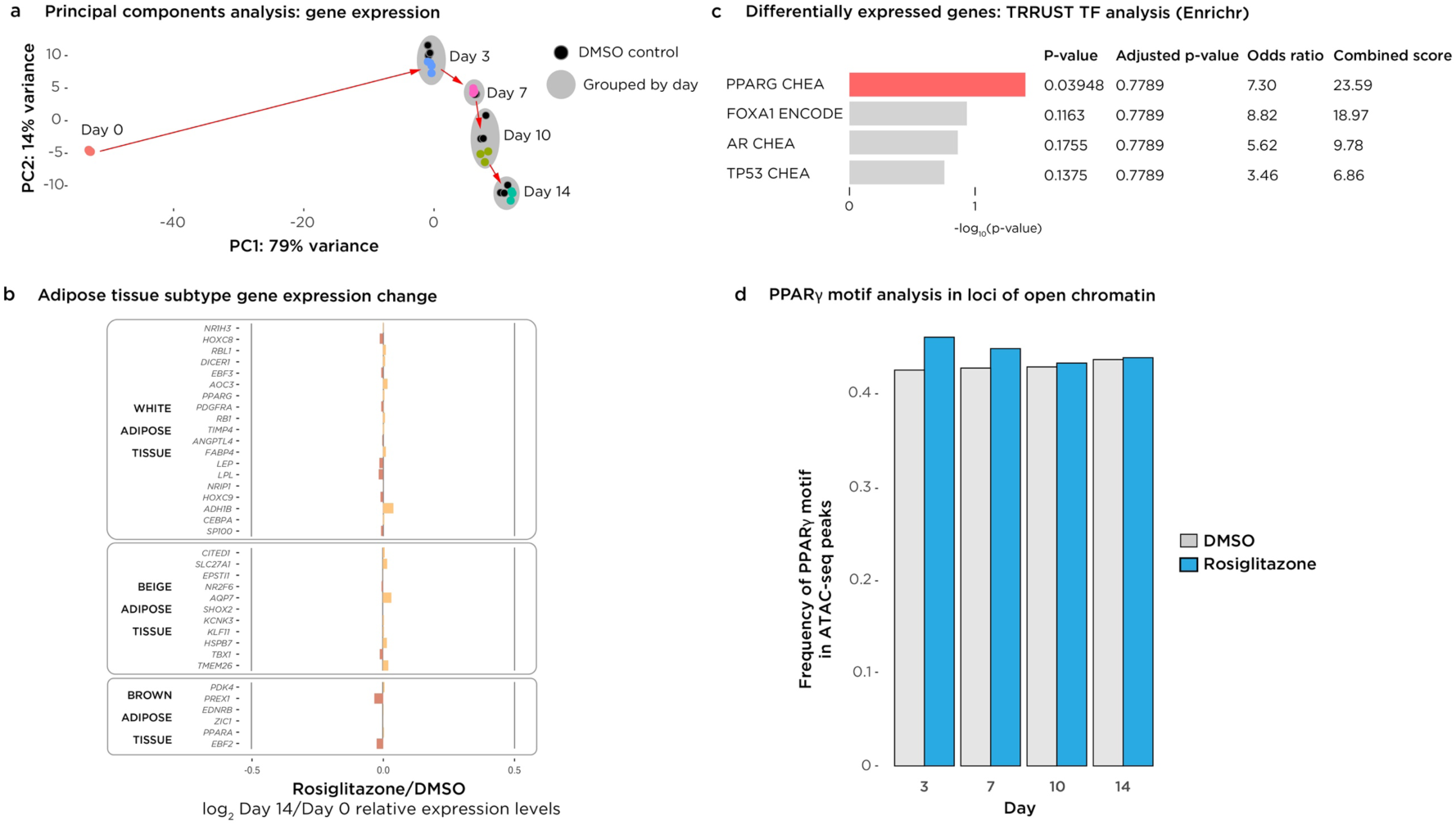
The PPARγ agonist Rosiglitazone has limited effects during early adipogenic differentiation. In (a) we use a principal components plot to represent the transcriptional variability at each timepoint, showing how the Rosiglitazone and DMSO control cells have near-identical expression patterns. In (b) we compare the day 14/day 0 expression patterns for Rosiglitazone exposure with the equivalent DMSO data, showing minimal changes and allowing us to conclude that the cells still produce white adipose tissue with Rosiglitazone exposure. In (c) we show the results of a ChIP Enrichment Analysis (ChEA) that reveals genes upregulated with Rosiglitazone exposure to be enriched for local binding of the PPARG TF. In (d) we quantified the proportion of peaks with the canonical PPARγ binding motif at each timepoint for the Rosiglitazone-exposed and control cells, showing a small enrichment at days 3 and 7.

ATAC-seq data also showed some evidence for PPARγ effects. In **Fig. 4d** we show the results of a targeted analysis, in which the peaks distinguishing the Rosiglitazone and DMSO control conditions were explored for the presence of the PPARγ binding site motif. The proportion of differential peaks with this motif was higher at days 3 and 7, supporting a model of Rosiglitazone acting early in adipogenesis at the chromatin level, followed by the transcriptional changes shown in **Fig. 4c** occurring subsequently at days 10 and 14.

### Tributyltin (TBT) exposure induces fatty acid synthesis genes in developing adipocytes

Our expectation from prior studies was that TBT exposure would be associated with PPARG:RXRA-mediated transcriptional and chromatin responses^3, 7, 8^. Our results indicate that the response to TBT is much more extensive and complex in differentiating MSCs. In **Fig. 5a** we show the PCA plot of gene expression variation over the time course. Whereas the effects of Rosiglitazone were minimal compared with the stage of differentiation, by Day 3 the TBT-exposed cells already have a markedly different pattern of transcriptional activity that persists through Day 14. This is reflected by modest changes in the expression levels of genes characteristic of white, beige and brown adipocytes, all less than a log_2_ magnitude of 0.4 (**Fig. 5b**), leading us to conclude that the differentiation continues to generate white adipose tissue. A summary heatmap of the genes with the greatest transcriptional changes is provided in **Supplementary Fig. 2**.

**Figure 5:**
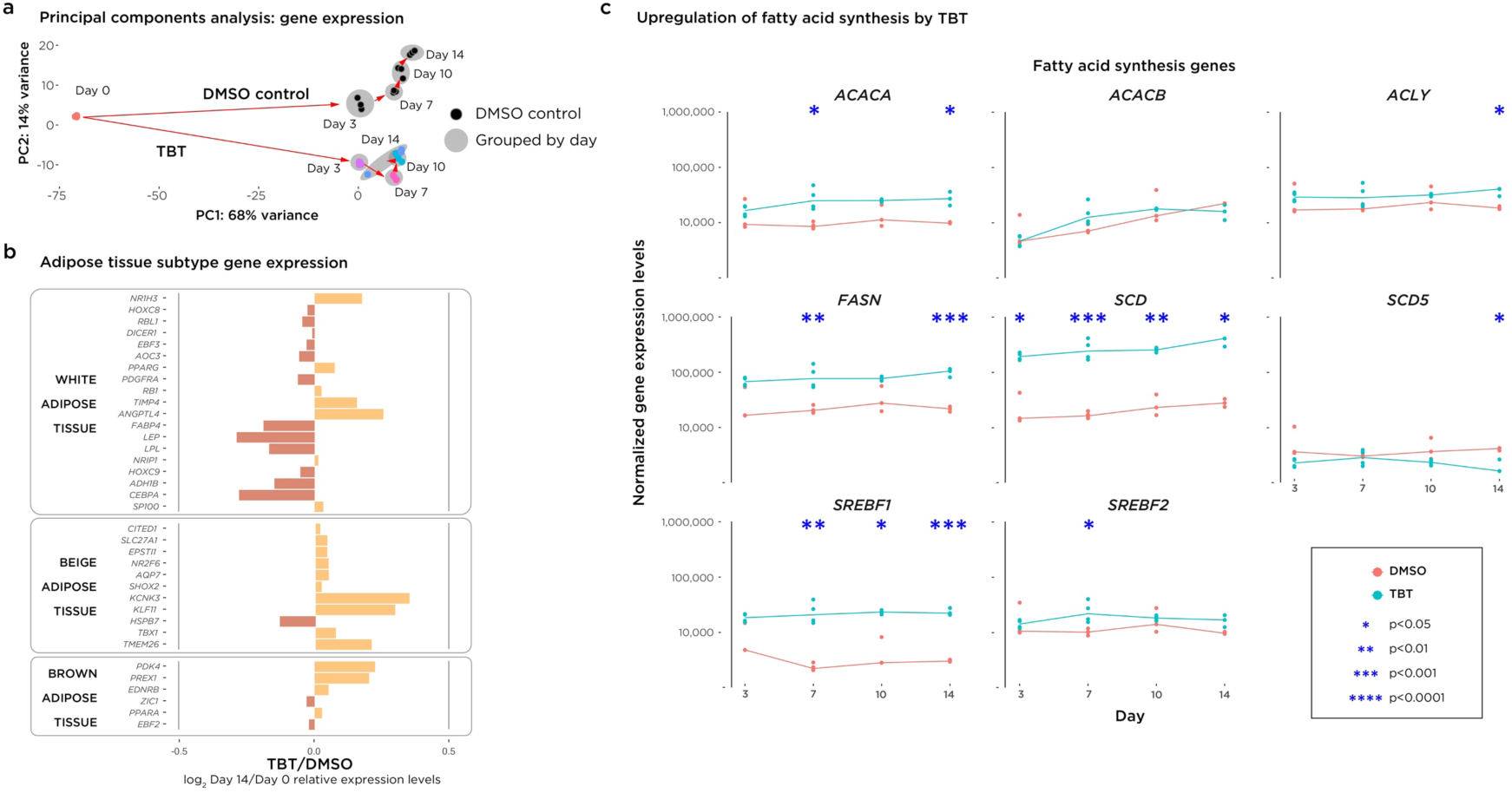
The transcriptional response to TBT exposure. In (a) the principal components plot shows that TBT induces striking differences in gene expression by day 3 that are sustained through day 14. In (b) we show that there are some modest changes of expression of the gene panels, but not enough uniformity or degree of change to indicate a change from white adipose tissue production. Panel (c) shows the panel of genes involved in fatty acid synthesis to be generally and significantly over-expressed with TBT exposure, implicating this as the metabolic process underlying the increased adipogenesis in these differentiating cells.

As our imaging cytometry studies showed that TBT causes an accumulation of intracellular neutral lipids (**Figure 1c**), we used the gene expression data to perform a focused exploration of the mechanism of this lipid accumulation. We hypothesized that this could be mediated by increased triglyceride or fatty acid synthesis, decreased fatty acid oxidation or lipolysis, or some combination of these processes. By comparing the expression levels of genes associated with each metabolic process^39–41^ between the TBT and DMSO control conditions, we were able to show a strong induction of genes involved in fatty acid synthesis^42^ (**Figure 5c**), with fewer changes in genes associated with the other metabolic processes (**Supplementary Fig. 3a-c**).

### Chromatin changes associated with TBT exposure

We note that while the transcriptional changes at Day 3 are marked (**Fig. 5a**), analysis of chromatin accessibility revealed only minimal changes, as shown in **Supplementary Table 5**. To test this unexpected observation further, we quantified the change in number of reads per peak (the unfiltered ‘concentration’ output from DiffBind), showing that the Day 3 changes are markedly less than those at later stages (**Supplementary Fig. 4**). We conclude that the major changes in expression at Day 3 are not dependent upon chromatin remodeling but are more likely to reflect cell signaling and transcription factor activation and inactivation acting at loci that are already accessible in the genome.

After the Day 3 time point there are abundant changes in the amplitude and locations of the ATAC-seq peaks. To understand whether there was a biological coherence to the genes targeted for these changes in chromatin accessibility, the genes at which these changes were occurring within ±1 kb of their transcription start site (TSS) at Day 10 (from **Supplementary Table 5**) were used to identify over-represented biological pathways. Those consistently enriched through differentiation were in two major categories, genes involved with the extracellular matrix, and genes encoding Rho, Rac1 and Cdc42 GTPases (**Fig. 6**).

**Figure 6:**
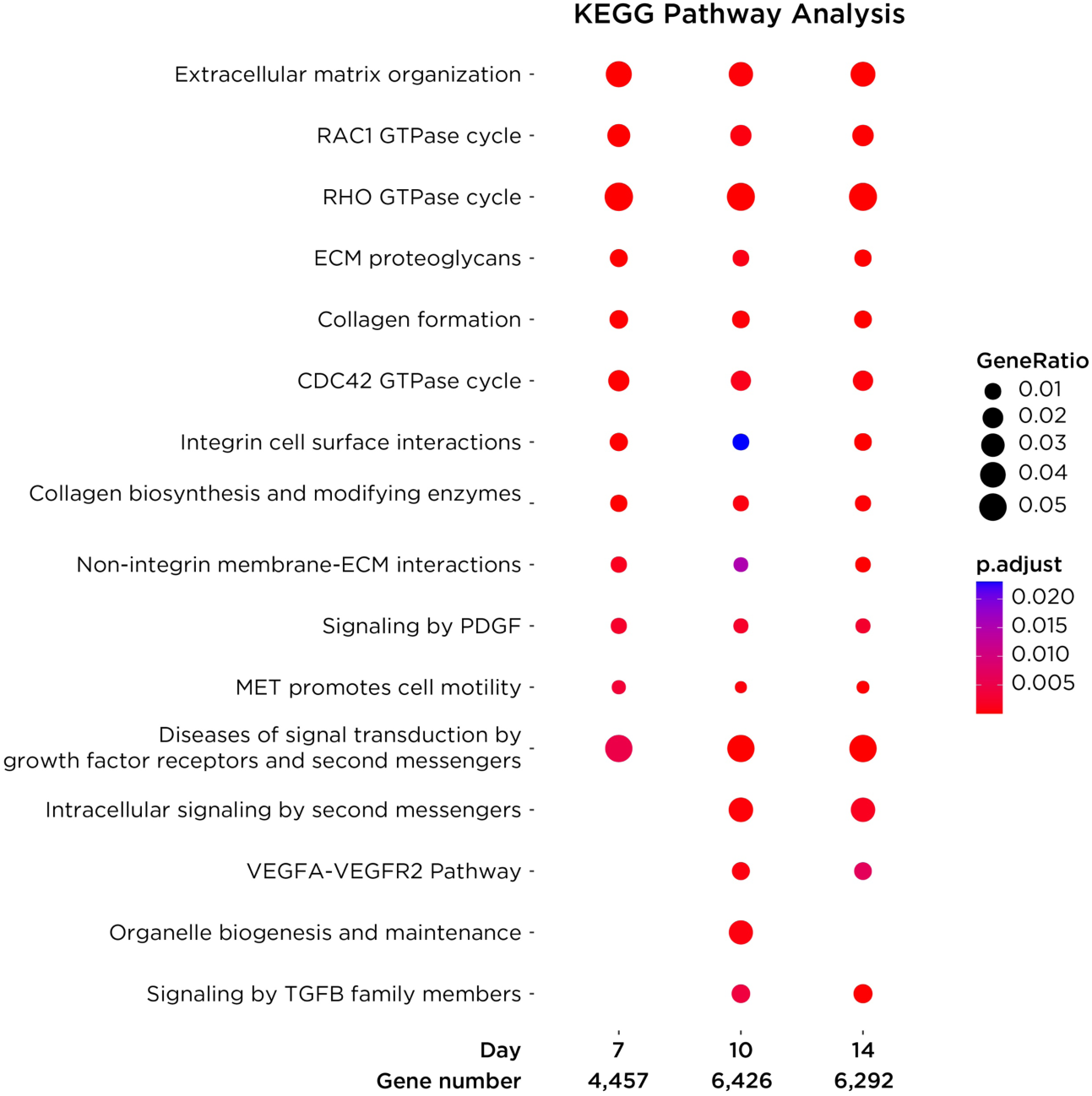
TBT-associated chromatin accessibility changes target the promoters of genes with specific properties. When the loci changing chromatin accessibility with TBT were linked to transcription start sites within 1 kb, the resulting gene list is enriched for Gene Ontology (GO) terms in two major groups: properties of the extracellular matrix, and Ras-related GTPases.

We then repeated the study represented by **Fig. 3**, identifying the TF binding motifs associated with the loci where the ATAC-seq peaks differed most substantially in TBT-exposed cells from the DMSO control condition. These results are represented in **Fig. 7** (with full results in **Supplementary Table 6**). The limited number of changes at Day 3 limit our analyses for this time point, but we see three groups of TFs emerging at the days 7, 10 and 14 time points. Those shown in black are the TFs that were found in **Fig. 3** to be expected at that time point, while those shown in blue are newly enriched with TBT exposure, and those in red and in parentheses were over-represented in normal differentiation and have become less frequent. The most consistent pattern throughout the time course is the inferred loss of binding of the TEAD and CEBP families of TFs, and a later increase in ETV family activity. CEBP TFs are long recognized to induce adipogenesis^27^, whereas TEAD TFs repress adipogenesis, at least in part through direct actions at the *PPARG* locus^29^.

**Figure 7:**
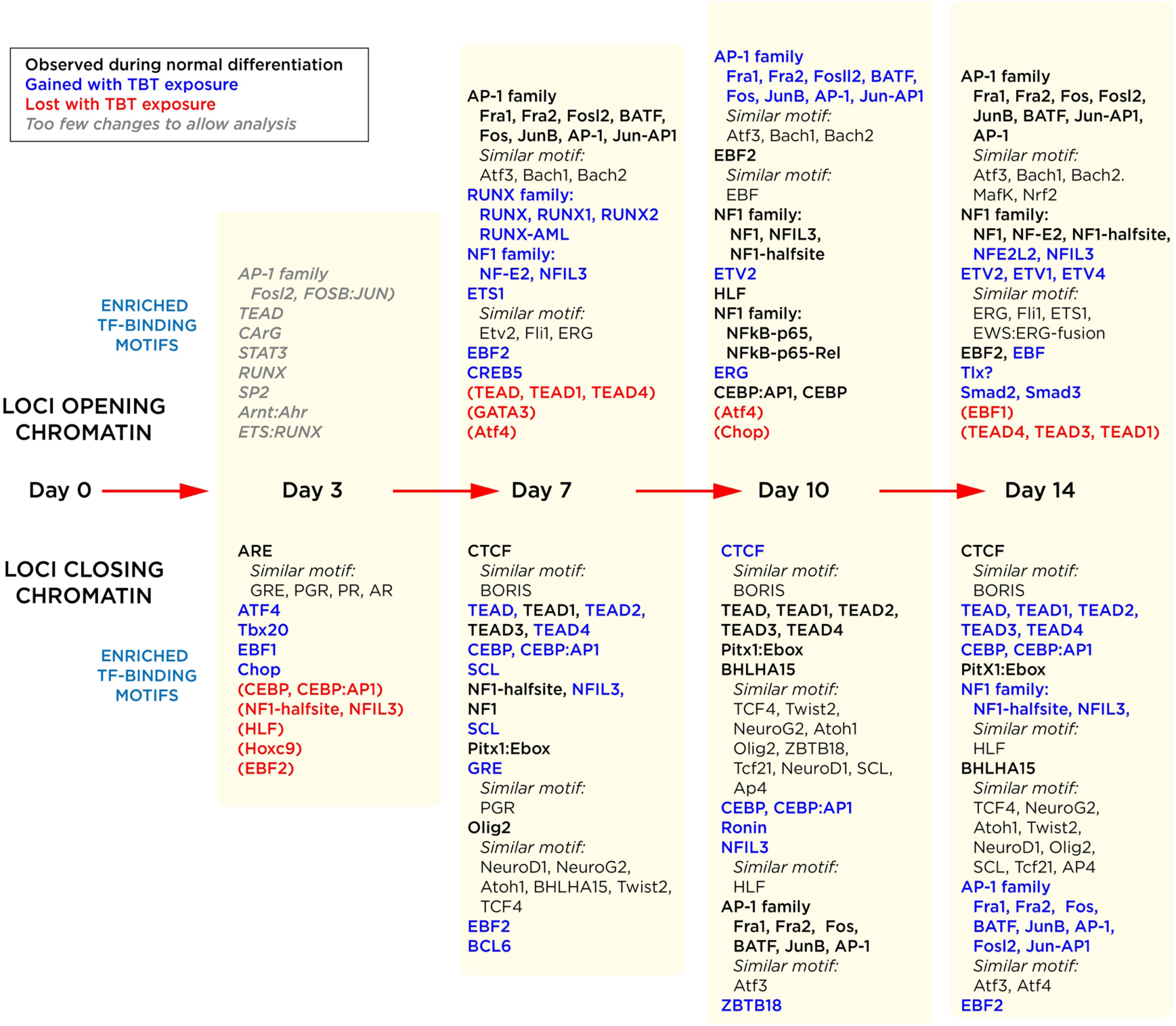
Changes of chromatin accessibility with TBT exposure implicate specific TFs. In this analysis the loci changing chromatin accessibility are comparted for the same timepoint between the TBT and control (DMSO) conditions. The day 3 changes were so few that we were unable to test loci gaining accessibility with TBT (upper), and the number of loci losing accessibility with TBT (lower) was so limited that the results at this time point should be interpreted with caution. The data from day 7 onwards are based on large numbers of loci and are more robust. The TF motifs that were already characteristic for these time points in normal differentiation are shown in black, with those newly gained in blue, and those no longer apparent in red and parentheses. Among the many changes, two stand out, the consistent loss of TEAD family TF motifs from day 7 on, and the loss of CEBP family TF motifs from day 10-14, as part of a broader group of regulatory changes. Despite attempting to include a broad representation of TF motifs, contrary to our prior expectation neither PPARG nor RXR motifs emerge from this analysis.

## DISCUSSION

Our studies confirm prior, groundbreaking work that showed progenitor cells to be altered by *in vitro* TBT exposure to increase adipogenesis^8^. In differentiating MSCs we separate the effects of increased cell number (hyperplasia) from increased lipid content per cell (hyperplasia), showing that TBT enhances both as part of its adipogenic effects. Our evidence supports the *in vitro* differentiation of the adipose-derived ASC52telo MSC line to white adipose tissue. By performing a time course approach involving multiple experimental replicates, we were able to develop a reference dataset for the gene expression and chromatin accessibility dynamics that occur during early adipogenesis. The most extensive changes in gene expression are those that occur immediately upon transferring the cells into adipogenesis-promoting tissue culture medium, with progressive changes thereafter that we could characterize in terms of trajectories of gene expression. We also were able to infer the identities of the TFs mediating these changes in chromatin accessibility by identifying over-represented short DNA sequence motifs in the loci of open chromatin, and linking them to known TF binding site motifs. The results obtained were reassuringly consistent with prior insights into the transcriptional regulation of adipogenic differentiation.

These temporal analyses of genome-wide transcriptional and chromatin studies have not been a feature of prior research into the effects of environmental obesogens on adipogenic differentiation. We were able to explore prior mechanistic models that were based on candidate mediators of adipogenesis, while allowing new, unbiased discoveries to emerge from our data. Our first analysis was focused on the PPARγ agonist Rosiglitazone. This drug has been shown to promote adipogenesis in an *in vitro* model of murine MSC differentiation^20, 43, 44^, which we were able to replicate in cells 21 days into adipogenic differentiation. Our molecular genomic studies focused on earlier events in MSC differentiation, with overall limited evidence for transcriptional or chromatin changes even at the highest Rosiglitazone doses. It is not possible to perform statistical testing on the few changes we observed, but we saw some indications of increased PPARγ binding to accessible chromatin earlier in the time course and increased expression of PPARγ target genes at days 7-14, suggesting that the Rosiglitazone was having modest effects on the differentiating cells. It has been previously proposed that TBT effects in murine MSCs in a similar *in vitro* system occur through a PPARγ-independent mechanism^45^, consistent with our findings.

Much more substantial changes in both gene expression and chromatin organization were associated with TBT exposure to the differentiating cells. As early as day 3 there were clear differences in gene expression, although this was not accompanied by correspondingly widespread alterations in chromatin accessibility. We were concerned that this chromatin accessibility observation was misleadingly due to a technical or analytical problem, but our further testing of the normalized read numbers in peaks confirmed the minimal differences in the days 0 to 3 comparisons, with manual scrutiny of genome browser representations of the data also consistent with the finding. We are therefore confident that these early transcriptional changes are occurring without a chromatin remodeling response.

We used the transcriptional information to gain insights into the cellular reprogramming occurring in the differentiating MSCs. While the genes in the panels used to distinguish white, beige and brown adipose tissues underwent some perturbation of their expression levels as part of the genome-wide alterations, neither the degree nor the consistency of changes appeared to indicate a shift from white adipose tissue production. When we compared the different potential mechanisms that could explain the increased lipid content per cell, the data most strongly supported a model of increased fatty acid synthesis as the primary metabolic effect of TBT exposure. Our transcription studies provided insights into the nature of the reprogramming by TBT of the adipogenic cells, yielding results consistent with the prior observation that TBT does not just generate more and larger adipocytes but a qualitatively different type of adipocyte^46^. This being the case, the possibility emerges that adipose tissue biopsies from human research participants could be profiled using these kinds of genome-wide assays to allow insights into whether their adipocytes reflect prior organotin exposure, a potential biomarker for the influence of environmental obesogens.

While TBT exposure is associated with a large number of transcriptional and chromatin organizational changes during adipogenesis, two distinct findings stand out that provide an insight into the possible primary mechanism for cellular reprogramming by TBT. We note in particular how ATAC-seq peak changes with TBT exposure that target the transcription start site region (±1 kb) predominantly involve genes encoding Rho, Rac and CDC42 GTPases, and the parallel observation that loci losing chromatin accessibility in TBT-exposed cells were enriched for TEAD TF binding site motifs. These Ras-related GTPases are involved in the maintenance of integrity of the actin cytoskeleton^47, 48^, and in fibroblasts can influence the organization of integrins as focal adhesions^49^ and thus the cell’s extracellular matrix interactions^50^. G protein-coupled receptor and Hippo signaling converge on the regulation of the transcriptional co-activators YAP and TAZ^51, 52^, with YAP entering the cell nucleus and binding to TEAD TFs to form a complex^53^ with enhanced transcriptional activation properties.

These observations point to a change in the cytoskeleton or extracellular matrix as a major feature of the cells exposed to TBT. Prior studies have tested the effect of TBT on the cytoskeleton, finding that rat thymocytes had loss of 80% of their F-actin content within 10 minutes of exposure, although at doses in excess of those used in our *in vitro* culture system^54^. A 2 minute *in vitro* exposure to human neutrophils at TBT concentrations comparable with those in this study caused significant reductions in F-actin content^55^, with other organotins showing comparable effects.

Our model for TBT-mediated influences on differentiating adipogenic cells is therefore as represented in **Fig. 8**, with a primary effect to depolymerize the F-actin of the cytoskeleton. The cell is then no longer able to sense physical influences, leading to the response by the Ras-related GTPases, the downregulation of YAP/TAZ transcriptional coregulators, and the loss of YAP-mediated TEAD family regulation of gene expression. As TEAD TFs repress adipogenesis^29^, the predicted effect of its loss would be obesogenic.

**Figure 8:**
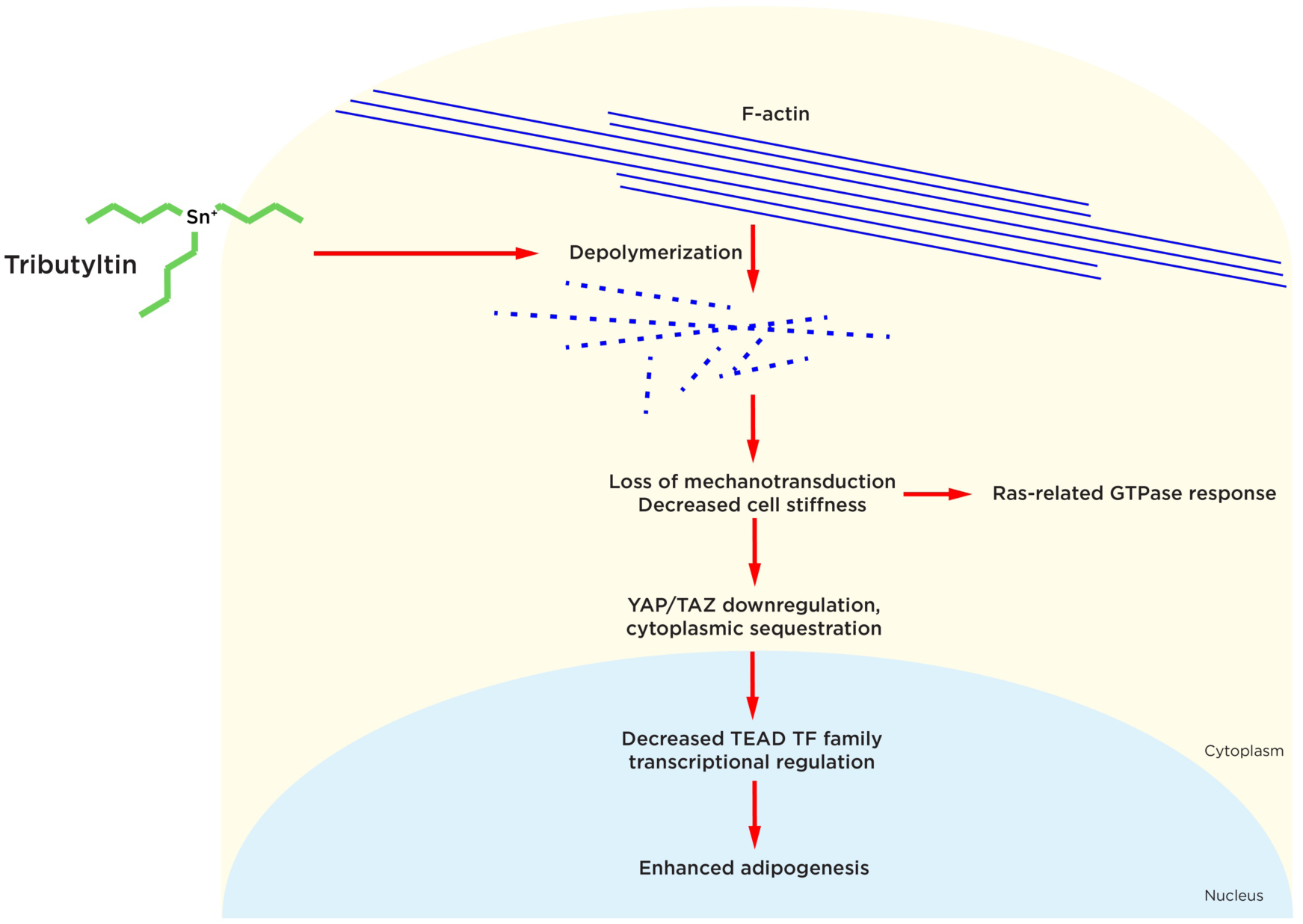
The model reconciling the molecular genomic findings of this study. Our data support a primary cellular effect by TBT to cause depolymerization of cytoskeletal F-actin, a known effect of TBT in mammalian cells. The resulting loss of mechanotransduction should lead to the response by the Ras-related GTPases that we observed, as well as the inactivation of the YAP transcriptional co-regulator, which then fails to activate TEAD TFs. As TEAD TFs function to suppress adipogenesis, the outcome should be one of increased adipogenesis.

This study complements and adds to prior investigations of the organotin group of environmental obesogens. We stress that while we are presenting a novel model for the adipogenic effects of TBT, our data do not exclude other well-studied models. While our genome-wide approaches are not highlighting PPARγ or RXR effects, this could be because they may act at a very small number of genomic loci, with effects that may be obscured when dealing with large numbers of findings. For example, our TF binding site motif analyses presented in **Figs. 3 and 7** necessarily had to be limited to those with the strongest significance. The limited effects of the PPARγ agonist Rosiglitazone may be in part due to our focus on earlier time points of adipogenic differentiation, with the possibility remaining that more extensive Rosiglitazone effects or PPARγ and RXR signatures in the genome might have been apparent if we had extended our studies further towards mature adipocyte differentiation. A major value of the study will be to prompt the further exploration of effects of organotins on the cytoskeleton and extracellular matrix, so that we can understand how to assay the effects of these and similar environmentally ubiquitous chemicals for their cellular effects with implications for human health.

## METHODS

### Mesenchymal Stem Cell (MSC) line culture and characterization

The cell line was originally derived from adipose tissue of a white female in 2009^17^. These human TERT-immortalized adipose tissue-derived mesenchymal stem cells (MSCs) were purchased from ATCC (ASC52telo (Catalog No. SCRC-4000); batch number 70003596). The cells were maintained following the ATCC-suggested protocol using Mesenchymal Stem Cell Basal Medium (ATCC Catalog No. PCS-500-303) with addition of the Mesenchymal Stem Cell Growth Kit for Adipose and Umbilical-derived MSCs - Low Serum supplement (ATCC Catalog No. PCS-500-040), and antibiotics (penicillin/streptomycin and G418).

The MSCs were differentiated into osteoblasts, chondrocytes, and adipocytes to validate multipotency. To differentiate cells into the adipogenic and osteogenic lineages, the MSCs were first transferred into StemXVivo Osteogenic/Adipogenic Base Media (‘base medium’ R&D Systems). To promote adipogenesis, Human/Mouse/Rat StemXVivo Adipogenic Supplement (Catalog No. # CCM011) was added to the base medium, for osteogenesis Human StemXVivo® Osteogenic Supplement (Catalog No. # CCM008) was used, while for chondrogenesis the cells were transferred into StemXVivo Chondrogenic Base Media CCM005 to which the Human/Mouse StemXVivo Chondrogenic Supplement (Catalog No. CCM006) was added. The seeding densities used were 2.1x10^4^/cm^2^ for adipogenesis, 4.2x10^4^/cm^2^ for osteogenesis, while chondrogenesis used 2.5x10^5^ clustered in a 15 ml conical tube. The cells were grown for 28 days following which they were tested for characteristics of each lineage.

For adipogenesis, in-well staining with Oil Red O was performed, while for osteogenesis Alizarin red S was used, and for chondrogenesis Alcian blue was used. Cells were removed from differentiation media, rinsed and fixed. Cells were stained following manufacturer’s protocol. Once stained cells were imaged and photographed to validate trilineage potential. Immunohistochemistry stains can be visualized in **Supplementary Fig. 1b**.

### Flow cytometry

The cells were tested for the presence of stem cell markers CD90, CD73, and CD105 and the absence of CD45 using the BD Bioscience hMSC analysis kit (Catalog No. 562245). Cells were detached from plate using trypsin and stained using the manufacturer’s recommended protocol. The FACS data are shown in **Supplementary Fig. 1a**.

### Chemical exposure conditions during adipogenic differentiation

Tributyltin Chloride (TBT) was purchased from Sigma Aldrich (Catalog No. T50202-500G). Rosiglitazone was purchased from Caymen Chemicals (Catalog No. 71740). The MSCs were seeded in the base medium at 2.1x10^4^/cm^2^, and allowed to grow (∼2 days) to >90% confluence. This defined day 0 of the differentiation. TBT and Rosiglitazone were diluted using dimethylsulphoxide (DMSO) as a vehicle, adding TBT (5, 50, 100 nM), or Rosiglitazone (10, 100, 500 nM) or a DMSO vehicle control (0.1% volume) to the differentiation medium starting at day 0. The concentrations of each chemical were based on previously published data looking at these dosages in 3T3-L1 and bone marrow derived MSCs^19, 44^. The differentiation medium consisted of the base medium plus the Human/Mouse/Rat StemXVivo Adipogenic Supplement. The culture was maintained using these conditions for up to 22 days.

### Oil Red O colorimetric assay

The Oil Red O assay was used to quantify neutral lipid accumulation in cells. The cells were rinsed with phosphate-buffered saline (PBS) and then stained in the culture well using the Adipogenesis Assay Kit (cell-based, Abcam Catalog No. ab133102). The dye was eluted using the kit’s elution buffer and quantified using a microplate reader at 490 nm wavelength. Empty wells were stained simultaneously as background controls.

### Celigo imaging and analysis

At day 8 of differentiation in the different exposure conditions, the wells were rinsed with PBS and then the cells were stained for neutral lipids using BODIPY 493/503 for 30 minutes at 37°C. The stained cells were then washed again with PBS and counterstained for nuclear imaging with Hoechst 33342 for 30 minutes at 37°C in the dark. The stained cells were then imaged using the Celigo Imaging Cytometer (Nexcelom). The green (BODIPY) and blue (Hoechst) intensities were quantified per well and compared with both the DMSO and empty control wells. The imaging cytometry primary data are provided in **Supplementary Table 1**.

### RNA-seq library preparation

The cells were differentiated in 6-well plates into adipocytes in the presence of TBT, Rosiglitazone, and DMSO (control) conditions for 3, 7, 10 and 14 days. RNA was isolated following washing of the cells using PBS and extraction with the Qiagen Micro RNeasy kit (Catalog No. 74004) following the manufacturer’s protocol. The cells were lysed in each well using the RLT buffer before transferring to microcentrifuge tubes to complete the protocol. The RNA concentrations were calculated using the Qubit 4. Bioanalyzer studies were used to measure the RNA integrity number (RIN) for each sample, using a cutoff of 8 for high-quality RNA. At least 3 samples from each condition and time point were sent to the New York Genome Center (NYGC) for sequencing. Their library preparation used the KAPA Stranded RNA-seq Kit, followed by 2 x 100 bp paired end sequencing on the NovaSeq to a depth of 30 million reads per library.

### RNA-sequencing analysis

The analytical methods used are available on our GitHub resource (see Code Availability below). Raw fasta files, normalized BAM files, and analyzed count matrices from library prep were downloaded from NYGC servers. Before downloading, 100 bp paired-end reads were trimmed to remove low quality base calls and Illumina universal adapters using Trim Galore! (version 0.6.5) with default parameters and then assessed using fastQC (version 0.11.4) and multiqc (version 1.10.1)^56^ The cleaned paired-end reads were aligned with STAR (version 2.5.2a) to GRCh38 genome, with Gencode v25 annotation. RNA abundance was measured by mapping with Bowtie2. BAM files were then used to create normalized counts matrices. Quality control was performed using Picard and RSeQC, and reads passing QC filters were aligned to annotated genes and quantified with featureCounts (v1.4.3-p1). Quality of raw data downloaded was validated by authors using FASTQC program (version 0.11.9)^57^ and Qualimap (version 2.2.1)^58^. All libraries passing QC metrics were used to create BAM files aligned to hg38 using the Bowtie aligner, then quantified with featureCounts (v1.4.3-p1). The raw featurecounts table was used for differential expression analysis. All downstream analysis of counts tables was performed in R (Version 4.2.0). Differential expression analysis was performed using DESeq2 version 1.36.0^59^ and gene set enrichment analysis was performed using clusterProfiler (version 4.0.5)^60^ and ReactomePA (version 1.40.0)^61^. Gene expression clustering was performed using K-means clustering with K set to 6, to identify patterns of gene expression alterations corresponding with changes during adipogenesis. Genes consistently upregulated with Rosiglitazone exposure from days 10 to 14 were identified (**Supplementary Table 7**) and used to perform Chip-X Enrichment Analysis (ChEA)^37^ with the Enrichr platform^38^, generating the results shown in **Fig. 4c.**

### ATAC-seq library preparation

The Assay for Transposase Accessible Chromatin (ATAC)-seq libraries were prepared using the published Omni-ATAC protocol^21^, with additional modifications to adjust for the high lipid concentration associated with adipocytes^62^. Cells grown concurrently with those used for the RNA-seq experiments were differentiated into adipocytes in 6-well plates using the same TBT, Rosiglitazone and DMSO exposures and the same 3, 7, 10 and 14 days duration. At each timepoint, wells were washed followed by the addition of 600 µl of 0.05% trypsin-EDTA was added to the cells for 7 minutes in the well at 37°C. Complete growth media (CGM) was then added to the well at 1.5x the volume of the trypsin-EDTA (900 µl) to stop the trypsin reaction. The cells were then gently scraped off the well then carefully pipetted up and down to break apart any cell clumps. The cell mixture was split in two and placed into prechilled 1.5 ml microcentrifuge tubes. A total of 20 µl of the cell mixture was added to 80 µl of PBS/Trypan Blue mixture for counting. 75,000 viable cells from each well were spun down at 650 RCF at 4°C for 5 minutes in a fixed angle centrifuge. The supernatant was removed and the Omni-ATAC protocol^21^ was performed, as modified by the Claussnitzer group^62^, increasing the IGEPAL concentration from 0.01% to 1% in Resuspension Buffer-I. A further modification was to double the transposase concentration during the transposition reaction in the same reaction volume. The libraries were quantified using the Qubit HS DNA kit (Life Technologies, Catalog No. Q32851). Library quality was assessed using 2% agarose gel electrophoresis and Bioanalyzer High Sensitivity DNA Assay tracings. A minimum of 3 libraries passing our quality assessments per condition and per time point were submitted to Novogene for 150 bp paired end sequencing using the Novaseq S4 flow cell, with an output of approximately 50 million reads per library.

### ATAC-seq data analysis

The analytical methods used are available on our GitHub resource (see Code Availability below). Sequencing data were downloaded from the Novogene platform. Fasta quality was checked using FastQC tool from Babraham Bioinformatics. Samples were then trimmed using TrimGalore and cutadapt from the Babraham Institute for paired end samples which removes the first 13 bp of Illumina standard adapters. Samples were then run again through FastQC^57^, and samples passing quality filter were aligned to the GRCh38 human genome assembly using Burrows-Wheeler Aligner (BWA) version 0.7.13 (bwa mem –M –t <N> hg38.idxbase <R2 FASTQ> <R1 FASTQ>)^63^ and then converted from SAM files with the parameter to remove low mapq >20 and any unmapped reads, after which the samples were sorted, indexed and prepared for further analysis. PCR duplicates were removed using samtools v 1.19 fixmate Mark Duplicates function in coordination with the markdup function. Flagstat QC was post duplicate removeal to ensure duplicate removal. Uniquely mapping reads were identified and then compared to the ENCODE list of highly represented regions^64^. Post-alignment quality checking was run through FastQC and ATACQC^65^ before calling peaks.

Peaks calling was performed using the MACS2^66^ Peak caller with the parameters -B –SPMR for paired end reads and a quality cut off of -q 0.01. Additionally, summits were called with each peak using the –call-summits command (macs2 callpeak --nomodel --nolambda -g 3e9 --keep-dup ’all’ --slocal 10000 -t <INPUT FILE> -n <OUTPUT PREFIX>)^67, 68^. Irreproducible Discovery Rates were defined for the overlapping peaks using the Nboley resource^69^. Peaks that passed the IDR threshold of 0.05 were retained and used to create consensus peak lists for each sample. The consensus peaks were used for all downstream analysis.

To determine the frequency of PPARγ motifs associated with peaks of differential accessibility, we used the MotifDb package in R with JASPAR2020 and TFBSTools packages to identify PPARγ-binding motifs at these loci. Differentially called peaks were identified by running ChipPeakAnno Overlaps on a set of consensus peaks per day that was generated by IDR analysis of narrow peak files using (module load idr/2.0.2/python.3.4.1- atlas-3.11.30\ python3 -c "import idr" \ idr ---samples). All replicates were run in paired analysis and then consensus sequences from each pair were also compared, forming a final consensus peak list. An IDR False Discovery Rate threshold was applied at each step, and any low quality peaks were removed. The consensus peaks in each group were read into R and then run through ChipPeakAnno (version 3.30.1), using its findOverlapsOfPeaks function. This produced a list of peaks that were condition-specific for each day. The results were then run through ChipPeakAnno’s makeVennDiagram function which uses hypergeometric testing to produce the final ranked list of peaks that are condition-specific.

The corresponding fasta sequences associated with each peak region were annotated using the getseq function from BSgenome (version 1.40.1). The MotifDB package was used to search these sequences for the PPARγ motif using the matchPWM function with an 80% minimum match score. The instances of PPARɣ motif occurrence were quantified in the peaks unique to Rosiglitazone and unique to DMSO for each day. The frequency of the PPARɣ motif was calculated by dividing the number of peaks with at least one PPARɣ hit by the total number of sequences peaks in each condition.

Differentially called peaks were defined using DiffBind (version 3.6.1)^70^ which used the previously created IDR consensus peaks and the cleansed aligned BAM files to generate a group of consensus peaks normalized to each sample’s background. Differentially opened chromatin regions were calculated DESeq2 analysis method in DIffBind. and Pathway analysis was performed with using clusterProfiler (version 4.0.5)^60^ and ReactomePA (version 1.40.0)^61^. Motif analysis was performed with HOMER (v4.11.9) to identify known motifs. Additional parameters and code can be found at the Github resource described in the **Code Availability** section.

## DATA AVAILABILITY

Genome sequencing data are available from the NCBI Gene Expression Omnibus (GEO) database:

https://www.ncbi.nlm.nih.gov/geo/query/acc.cgi?acc=GSE207040

## CODE AVAILABILITY

All scripts for analysis have been placed on our Github resource:

https://github.com/tthompson29/Tributyltin_Adipogenic_Genomic_and_Transcriptomic_Analyses

## Supporting information

aSupplementary Figure 1

## ACKNOWLEDGEMENTS

The authors thank Drs. Samantha Laber and Melina Claussnitzer (Broad Institute) for sharing details on their optimized experimental protocols. We thank Einstein colleagues Dr. Masako Suzuki, Reanna Doña-Termine, Marliette Matos Rodriguez, Jacob Stauber and Daniel Borger for their help with experimental design, assays, and data analyses. TVT was supported by the Einstein Medical Student Training Program (NIH T32 GM007288) and the NIH (NIEHS) award F31 ES031480. Einstein core facilities involved were the High-Performance Computing Core, the Epigenomics Shared Facility, and the Genomics Core Facility, with support from the Albert Einstein Cancer Center (P30CA013330) and the Center for Epigenomics. The work was supported in part by R01 AG057422 to JMG.

## AUTHOR CONTRIBUTIONS

TVT and JMG designed the experiments, interpreted results, and wrote the manuscript. TVT performed experiments, processed and analyzed data. TVT and JMG created figures.

## MATERIALS & CORRESPONDENCE

Materials requests and correspondence should be addressed to John Greally, Center for Epigenomics and Department of Genetics, Albert Einstein College of Medicine, Bronx, NY, 10461, USA.

