## Supplementary figures and images for "Molecular genomic studies of the obesogenic effects of tributyltin during adipogenic differentiation implicate a primary role for cytoskeletal damage"

### aSupplementary Figure 1

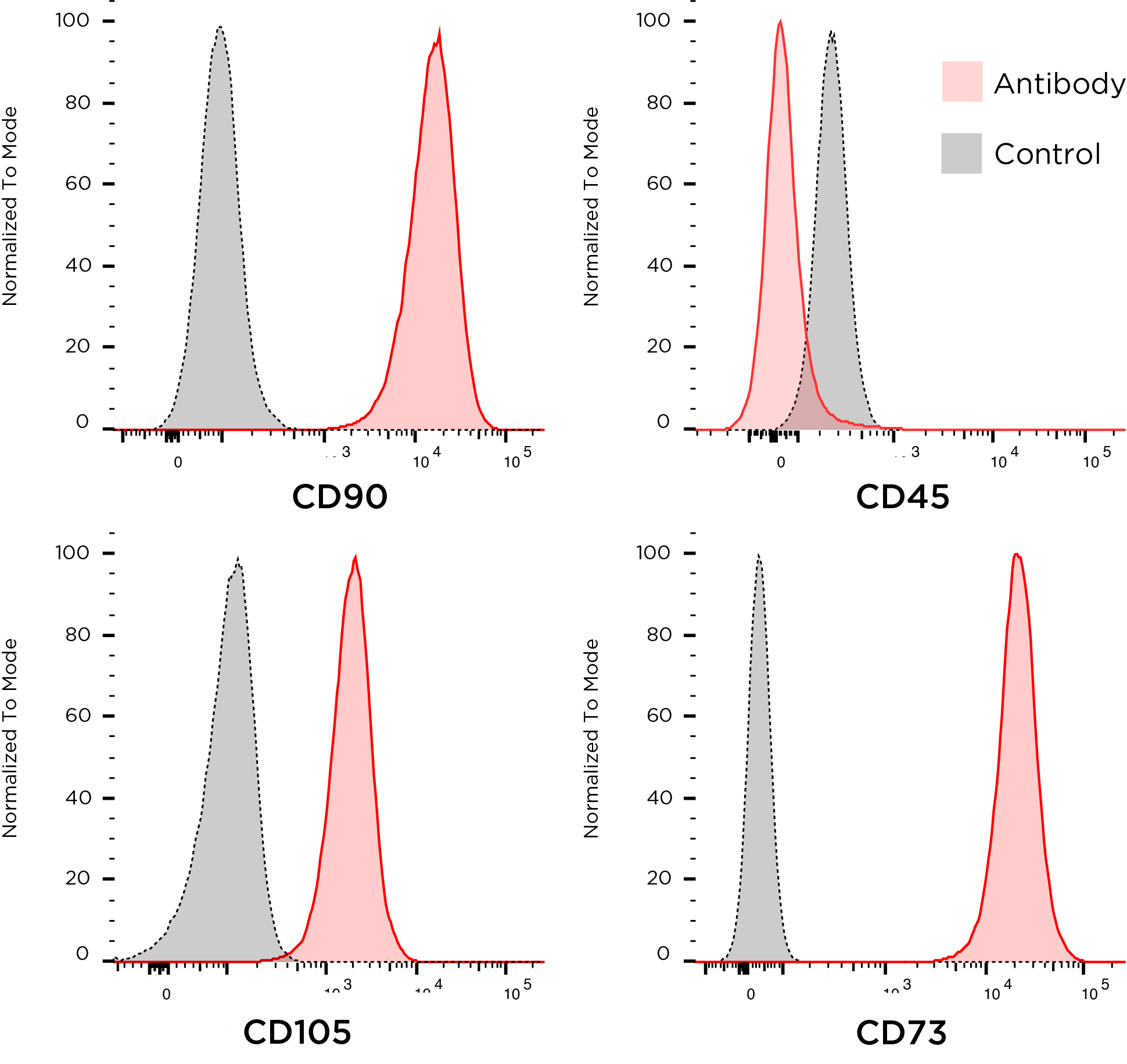

### Supplementary Figure 1b

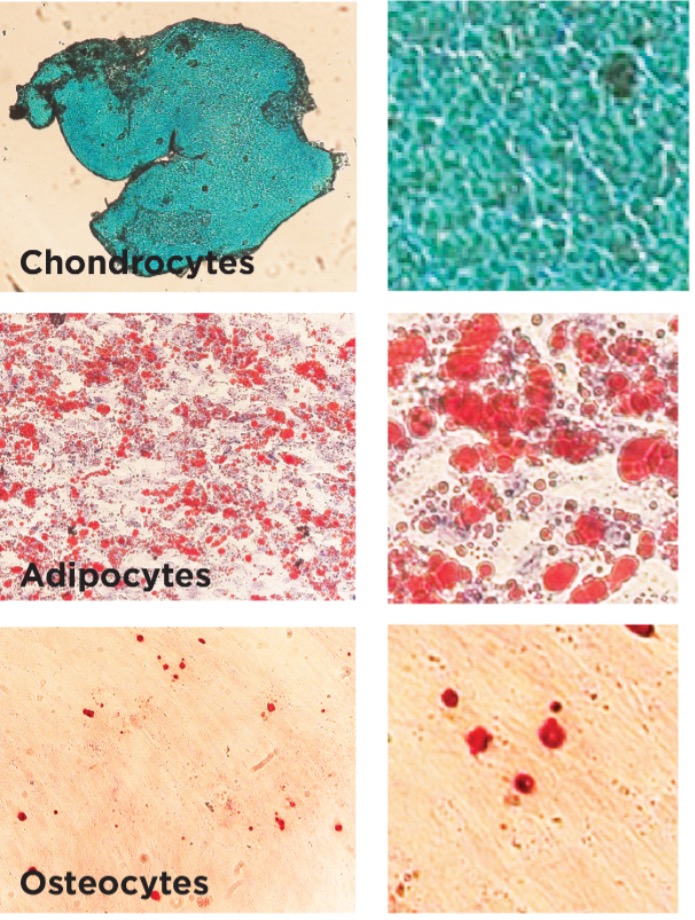

### Supplementary Figure 2

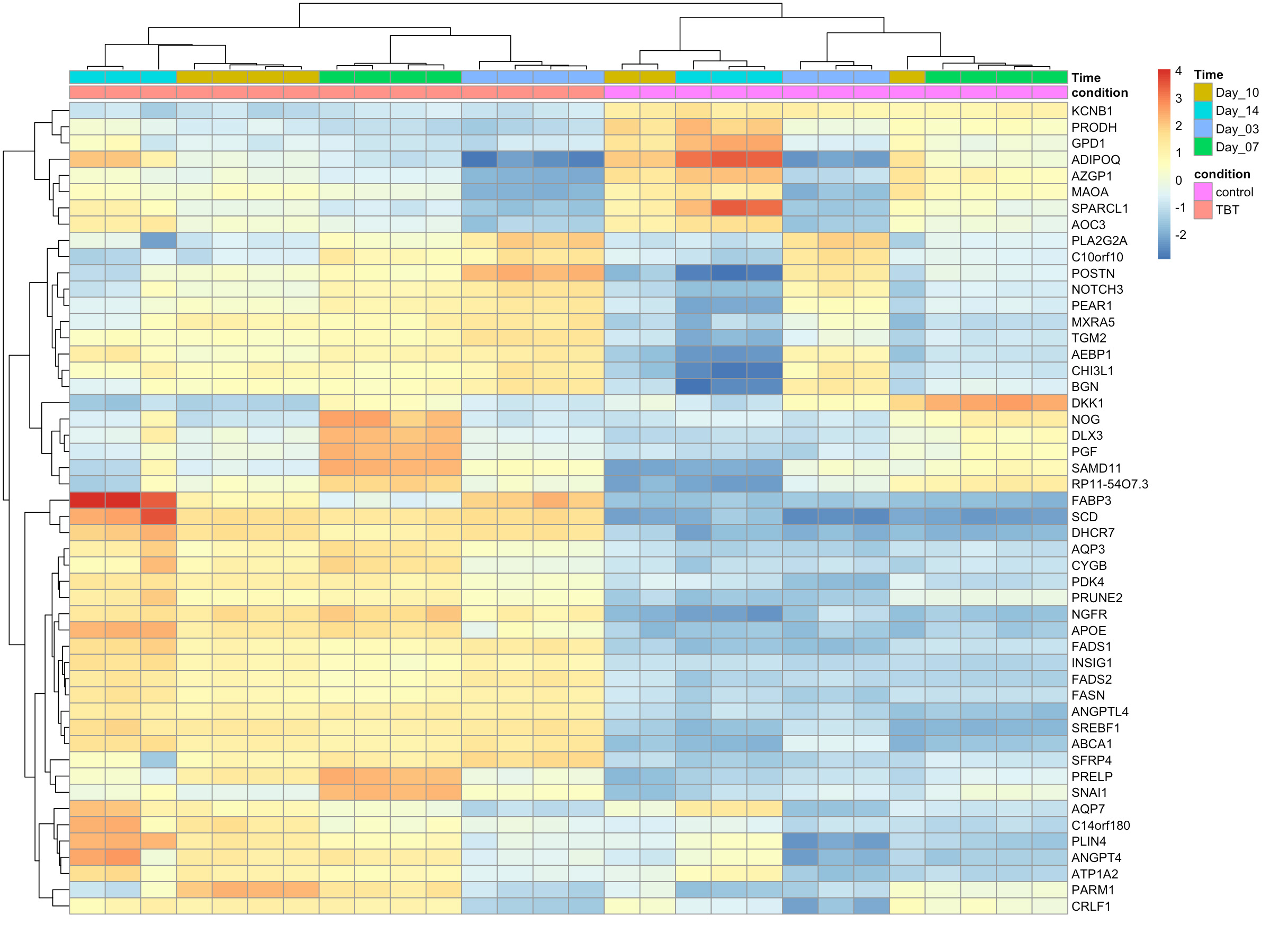

### Supplementary Figure 3a

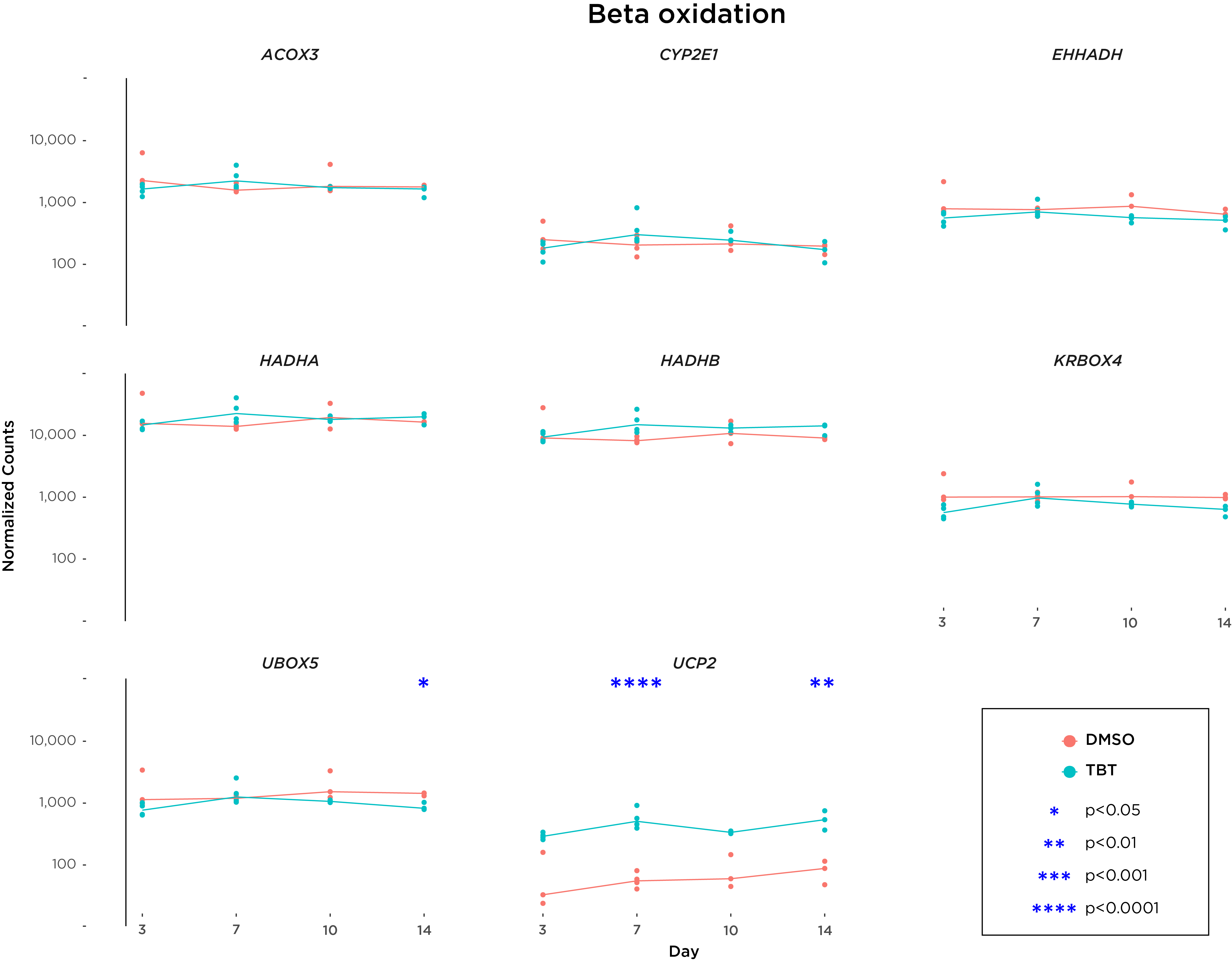

### Supplementary Figure 3b

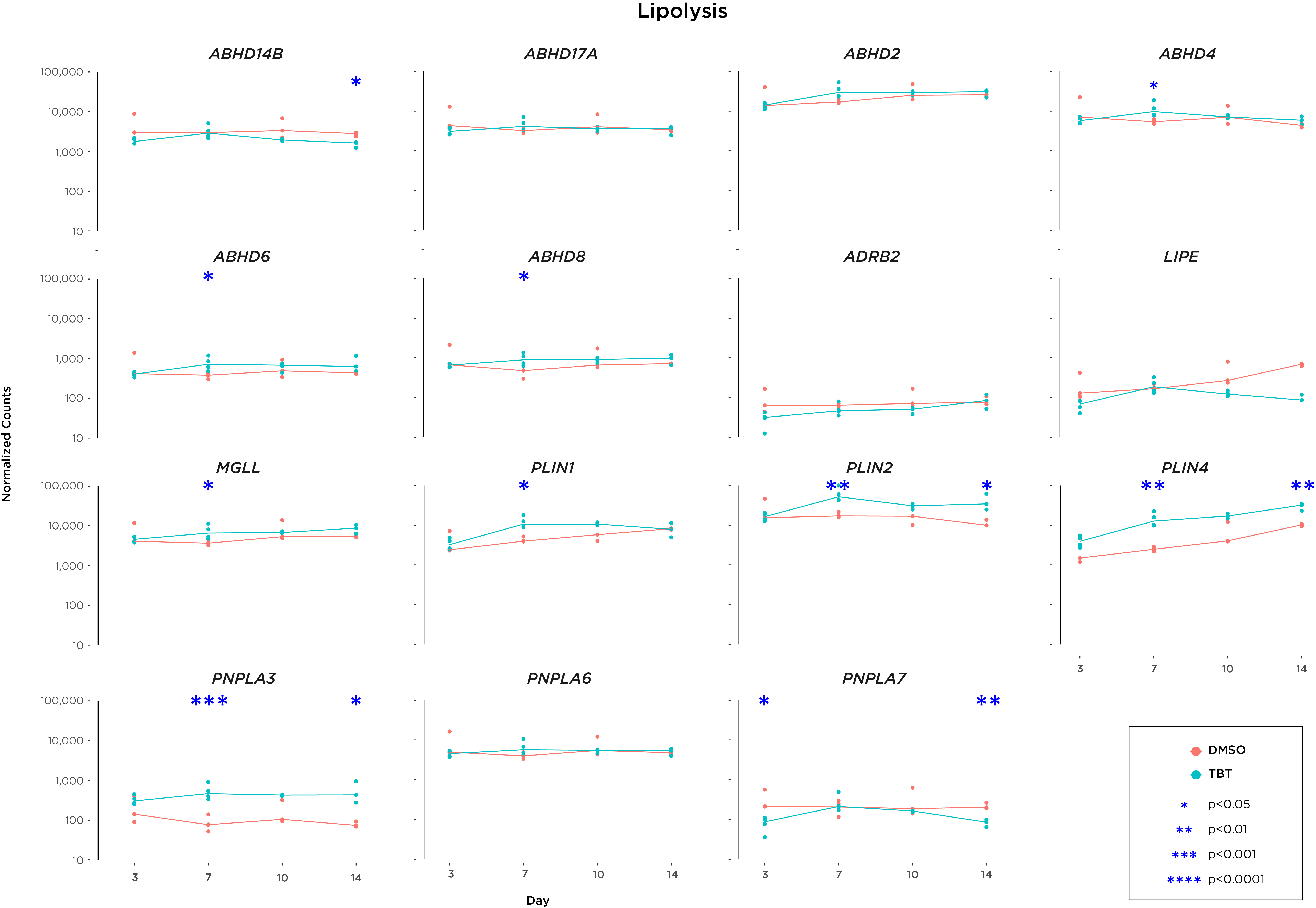

### Supplementary Figure 3c

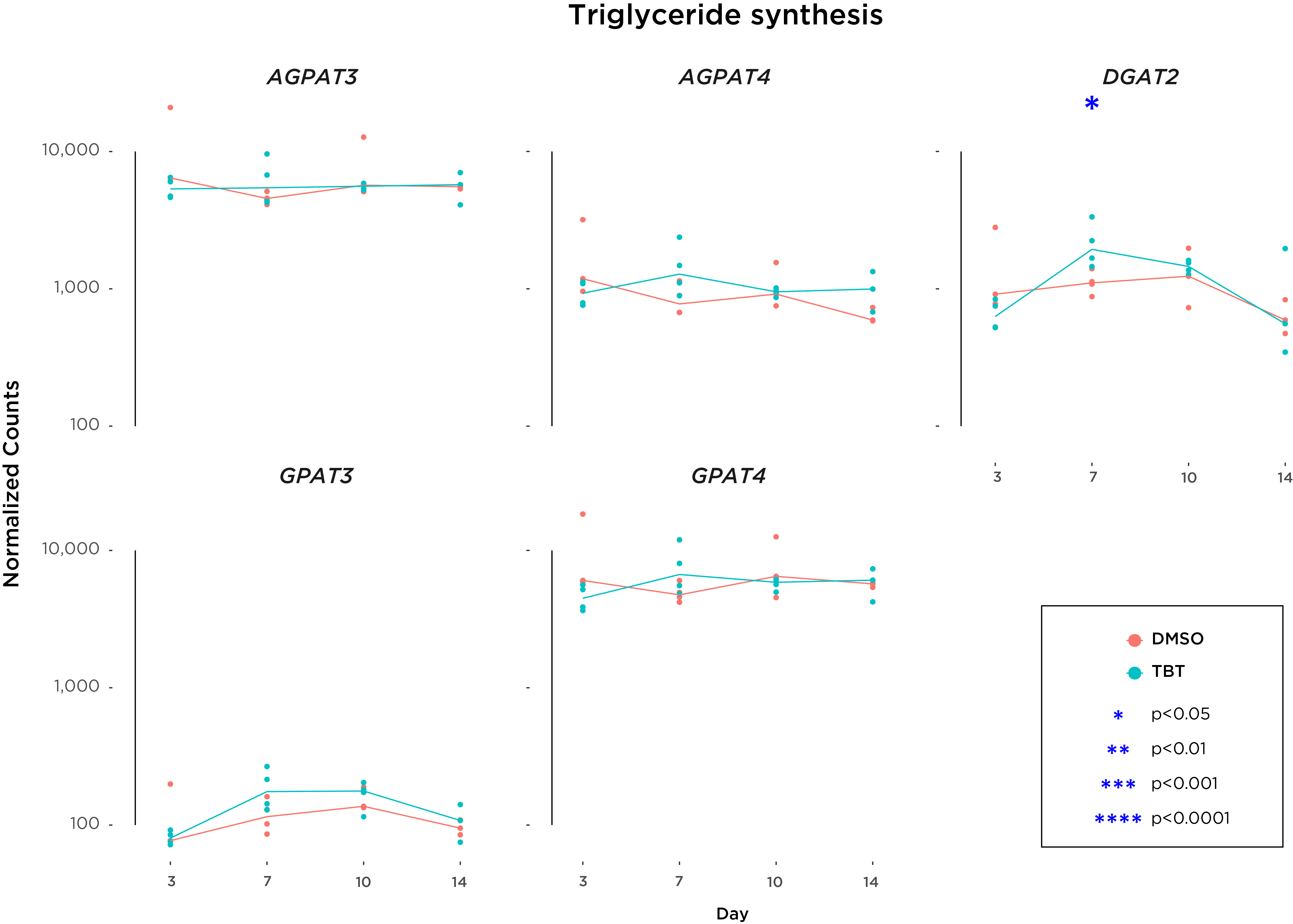

### Supplementary Figure 4

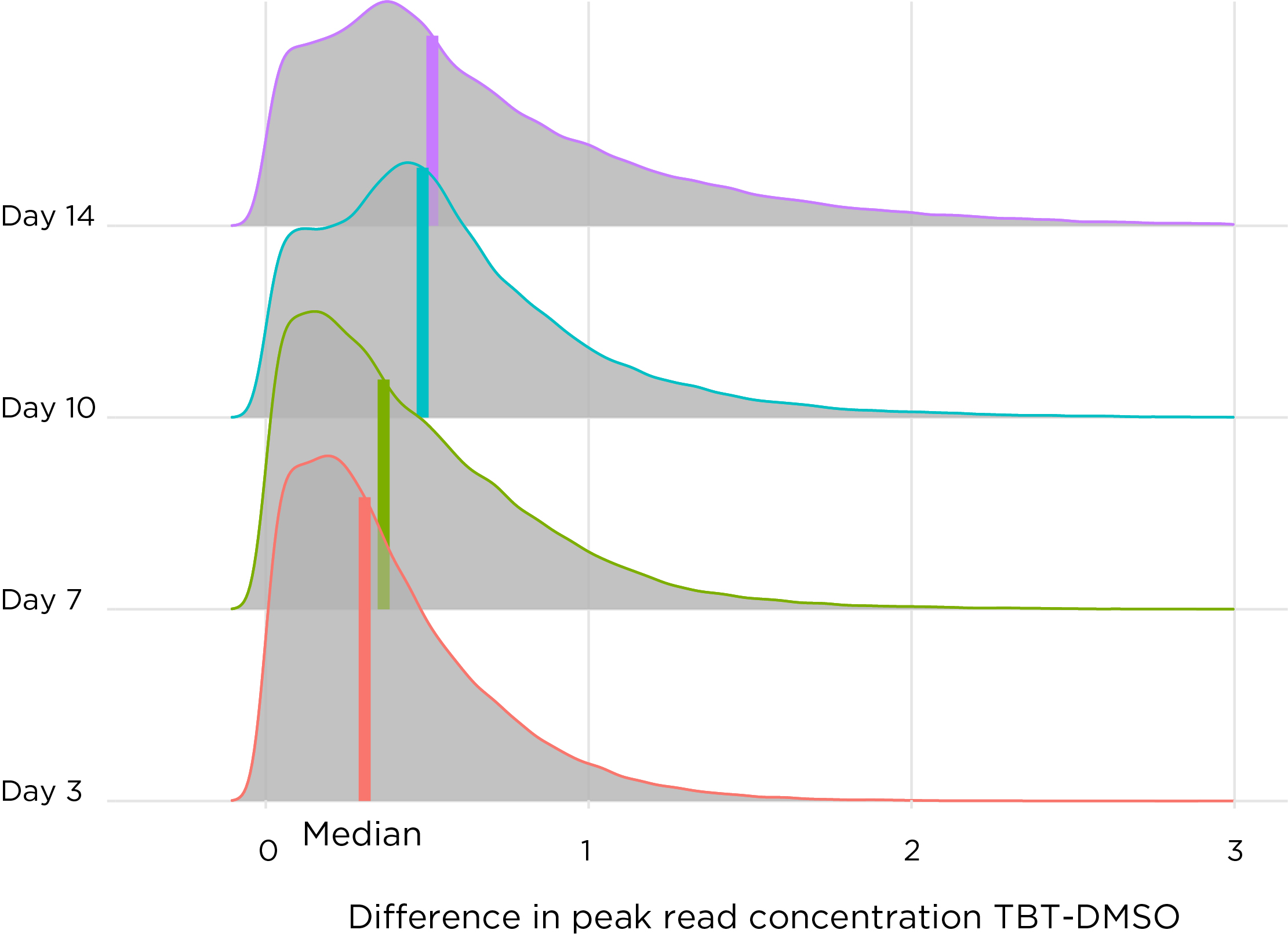
